# Reliability and precision of thermal imaging measurements to study animal behaviour and welfare

**DOI:** 10.1101/2025.07.31.668027

**Authors:** Debottam Bhattacharjee, Marianne A. Mason, Alan G. McElligott

## Abstract

The use of infrared thermal imaging has become increasingly popular in animal behaviour, health, and welfare research over the last decade. Yet, there is a lack of consensus regarding how this technique should be best applied when measuring peripheral temperatures in animals, including which regions of interest to favour. This fundamental issue necessitates checking the reliability and precision of thermal imaging data when taking repeated measurements, both over short and relatively long time windows. Using goats (*Capra hircus*) as a model, we investigated two subcategories of reliability, short-term repeatability (measurements taken in the same session) and reproducibility (over multiple sessions), as well as the precision of surface temperatures in two facial regions. We collected data from 20 goats over five measurement sessions over consecutive days. During each session, five frames were collected from approximately one-minute-long videos. From each video, we extracted the mean, maximum, and minimum surface temperatures from the left eye, right eye, and nose tip. To calculate repeatability, we compared temperature variation attributed to differences between goats against total variation in surface temperatures measured in a single session. We defined precision as the temperature deviation within which the mean temperature measured from one to five thermal images was expected to fall in relation to the mean of five image replicates 95% of the time. Reproducibility was investigated by comparing variation attributed to differences in temperature between measurement sessions against total variation in surface temperatures. Our results revealed that repeatability and precision of mean and maximum temperatures across five repeated measurements were high for all facial regions, with between 93.50% and 99.81% of total temperature variation attributable to the individual goat tested. Conversely, minimum temperatures were more variable, less repeatable, and less precise. For reproducibility, measurement sessions accounted for a high proportion of variation in nasal temperatures (74.61-85.85%), and a lower, but substantial proportion of eye temperature variation (49.59-67.01%). We conclude that mean and maximum thermal measures show promise for quantifying nasal and eye temperatures in the short term. However, surface temperature measured across several days was not readily comparable, highlighting the importance of considering ambient conditions in thermal imaging research. Overall, this study provides valuable insights into the appropriate use of thermal imaging in goats and, more broadly, animal behaviour and welfare research.

## 1. INTRODUCTION

Infrared thermal imaging is an approach increasing in prominence within the fields of animal behaviour, physiology, welfare, and veterinary research. Applications of this technique are based on the principle that all terrestrial objects with an absolute temperature exceeding zero kelvin generate radiant heat in the infrared region of the electromagnetic spectrum (Knízková et al., 2007; Speakman & Ward, 1998). Levels of infrared radiation emitted by objects can be detected via thermal imaging and used to generate a visual representation, allowing users to observe and quantify minute spatial and temporal disruptions in an object’s surface temperature (Knízková et al., 2007). When the object of interest is an endothermic animal, local fluctuations in blood flow, metabolic activity, tissue conductivity, and environmental heat exchange create a dynamic network of graded temperature zones occurring across the periphery of the skin (McCafferty et al., 2015; Godyń et al., 2019). By concentrating measurements on particular regions of interest (ROIs), generally areas where fur is thinner or absent, such as the eyes or nose, it is possible to investigate differences in skin temperature both between and within individuals, over time (Tattersall, 2016). Thus, these ROIs can help identify the internal physiological and emotional processes. For instance, diversion of blood away from peripheral blood vessels caused by stress acts to infuse core muscles and organs required in ‘fight or flight’ responses and redistributes heat, causing a drop in peripheral temperature. This redistribution of body heat during emotional experiences has been demonstrated in positive (Proctor & Carder, 2015; 2016; Tamioso et al., 2017) as well as negative contexts (Stewart et al., 2008a; 2008b; Herborn et al., 2015; 2018; Bhattacharjee et al., 2024; Ramirez Montes de Oca et al., 2024). Further, skin temperature changes caused by local increases in metabolism and blood flow can be indicative of inflammation in underlying tissues or differences in reproductive receptivity (Kominsky et al., 2010; Byrne et al., 2019; LokeshBabu et al., 2018; Façanha et al., 2018; Mota-Rojas et al., 2021). While the non-invasive application of thermal imaging is advantageous (Tattersall, 2016), the sensitivity of peripheral temperature and thermal imaging itself to a suite of different factors can make this technology difficult to apply with precision in practice.

Animal surface temperatures are affected by several factors. Endogenous factors, such as individual, breed, sex, age, level of physical activity, and skin and coat characteristics, can lead to variations in surface temperatures (thickness and colour; Bartolomé et al., 2013; Rizzo et al., 2017; Jørgensen et al., 2020; Jansson et al., 2021; Mota-Rojas et al., 2021). Environmental conditions, such as ambient temperature, humidity, and wind speed, also affect peripheral temperatures, either directly or via internal, homeostatic mechanisms (Church et al., 2014; Jansson et al., 2021). Further extraneous factors affecting accuracy of temperature readings include distance, or angle of the subject in relation to the camera, the camera model, the ROI used, its size, and the particular measure chosen, i.e., mean, maximum, or minimum temperature (Church et al., 2014; Howell et al., 2020; Ijichi et al., 2020; Playà-Montmany & Tattersall, 2021; Uddin et al., 2020; Kim & Cho, 2021). Particularly in measuring animal emotional responses, there is a distinct lack of consistency across investigations regarding ROIs chosen and specific temperature measures favoured. These have included, mean and minimum nasal temperatures (Proctor & Carder, 2015; 2016; Kano et al., 2016; Brügger et al., 2021; Bhattacharjee et al. 2024), maximum eye temperatures (Stewart et al., 2008a; 2008b; Bartolomé et al., 2019), temperatures in specific eye regions (e.g., lacrimal caruncle: Dai et al., 2015) and other regions, such as maximum temperatures of a chicken’s comb and wattle (*Gallus gallus domesticus*; Herborn et al., 2015) and a pig’s rear (*Sus scrofa domesticus*; Boileau et al., 2019). Thus, it is important to assess the reliability and precision of thermal imaging measures for best practices in animal behaviour and emotion research.

To assess reliability, the degree of similarity between repeated measures taken from the same ROI and the subject must be determined (Bartlett & Frost, 2008; Fernández-Cuevas et al., 2015). In humans, high levels of repeatability or repeated measurements taken under identical conditions, and reproducibility or measurements taken in variable conditions, i.e., the two subcategories of reliability (cf. Bartlett & Frost, 2008), have been found in a medical setting (Ammer, 2008; Petrova et al., 2018). By comparison, ambient conditions for animals can be more variable, e.g., in farms, captivity, or even open environmental conditions (Cilulko et al., 2013). Additionally, animals can be less compliant than humans, making obtaining accurate thermal imaging measurements in animals challenging. Attempts to investigate the reliability of thermal imaging measures have been made in cows (*Bos taurus*; Byrne et al., 2017; Scoley et al., 2019), sheep (*Ovis aries*; Byrne et al., 2019), and horses (*Equus caballus*; Howell et al., 2020). Yet, to our knowledge, no similar efforts have been made in goats (*Capra hircus*). Although applications of thermal imaging technology have been explored across a few contexts in goats (e.g., Façanha et al., 2018; Bartolomé et al., 2019; Giannetto et al., 2020), at least compared to other domesticated species (e.g., horses and cattle: reviews by Soroko & Howell, 2018; Mota-Rojas et al., 2021, respectively), it remains a relatively untapped resource in goat welfare and veterinary research (Mason et al., 2024a). Given the contribution of goats to emotion research (Briefer et al., 2015a, 2015b; Baciadonna et al., 2019; Mason et al., 2024b, 2025), identifying suitable ROIs and measures to quantify surface temperature reliably will not only be valuable for further goat research but can considerably advance the methodological aspects of thermal imaging research.

Although less commonly employed, video analysis offers advantages over extracting temperature data from a series of still images collected manually using a handheld device (McManus et al., 2022). Video recording substantially increases the number of measurement frames that researchers can collect in a short time frame. This, in turn, enhances the precision of animal surface temperature estimates (Byrne et al., 2017; Scoley et al., 2019; McManus et al., 2022), as well as enabling a finer-grained analysis of skin temperature changes over time (e.g., for measuring respiration rate: Stewart et al., 2017; Jorquera-Chavez et al., 2019; and time courses of emotional responses: Stewart et al., 2008a; 2008b; Herborn et al., 2015). Furthermore, given that video footage can be captured from fixed positions, i.e., next to objects that animals regularly interact with, such as feeders and automated milking systems (Hoffmann et al., 2013), it minimises the need for human proximity. This latter point is important as the presence of experimenters may affect animal physiological responses, potentially undermining the validity and generality of results obtained (Moe et al., 2017; Cannas et al., 2018). However, attempts to assess reliability and other practical aspects of this data collection method remain limited (Hoffmann et al., 2013; Cuthbertson et al., 2019; Jorquera-Chavez et al., 2019). Therefore, the objective of the current investigation was to evaluate short-term repeatability (measurements taken within a single session) and precision, as well as reproducibility (over five consecutive days) of goat mean, maximum, and minimum temperature measured in the eyes and nasal region, extracted from thermal imaging videos.

## 2. MATERIALS & METHODS

### 2.1. Ethics Statement

All animal care and testing protocols were in line with ASAB guidelines for the use of animals in research (Bee et al., 2021). Further approval was granted through an ethical amendment (03.21) made to the project ‘goat perception of human cues’ (Ref. LSC 19/ 280) by the University of Roehampton’s Life Sciences Ethics Committee. All procedures were non-invasive, goats were kept unrestrained and tested in groups to avoid social isolation, showing no obvious signs of stress during experimental trials.

### 2.2. Study Site & Sample Population

Experiments were conducted between 5^th^ and 23^rd^ July 2021 at Buttercups Sanctuary for Goats in Kent, UK (51°13’15.7“N 0°33’05.1”E; http://www.buttercups.org.uk/). During this period, the sanctuary was open to the public. It featured a large outdoor paddock to which goats had access during the daytime, before being housed individually or in small groups within a large stable complex at night (mean pen size = 3.5m^2^). Goats had hay, grass, and water available *ad libitum* throughout the day, and were supplemented with commercial concentrate contingent on age and condition. We selected subjects based on their ease of handling and, as goats were tested in groups of three, on whether they showed evidence of affiliative relationships with two other goats as indicated by staff at the sanctuary (to minimise instances of aggression). Our final sample size comprised 20 goats (10 intact females and 10 castrated males) of various breeds and ages (see Table S1).

### 2.3. Experimental Set-up

Experiments were carried out within a familiar stable that goats could freely access during the day. We kept the windows closed to minimise drafts and avoid direct sunlight falling on the subjects. Both the windows and two skylights directly above pens A, B, and C were covered (Figure 1). To improve consistency in imaging conditions between subjects and over repeated measurements, trials mainly took place within pen B, with other members of the subject’s group being placed in adjacent pens A and C. This was not always possible, and for two groups, some group members were placed in pen D to prevent agonistic interactions with the subject. Distractions from surrounding objects also meant two subjects were imaged in pen A so that their attention could be successfully directed towards the camera. Pens featured a hayrack to encourage subjects to spend time at the front of the pen (hay availability shown to elicit minimal emotional arousal in goats at the same study site: Briefer et al., 2015b). Water was available in pens A, C, and D, but not in B, to reduce the potential effect of the presence of moisture on the muzzle on goat nasal temperatures.

**Figure 1.**
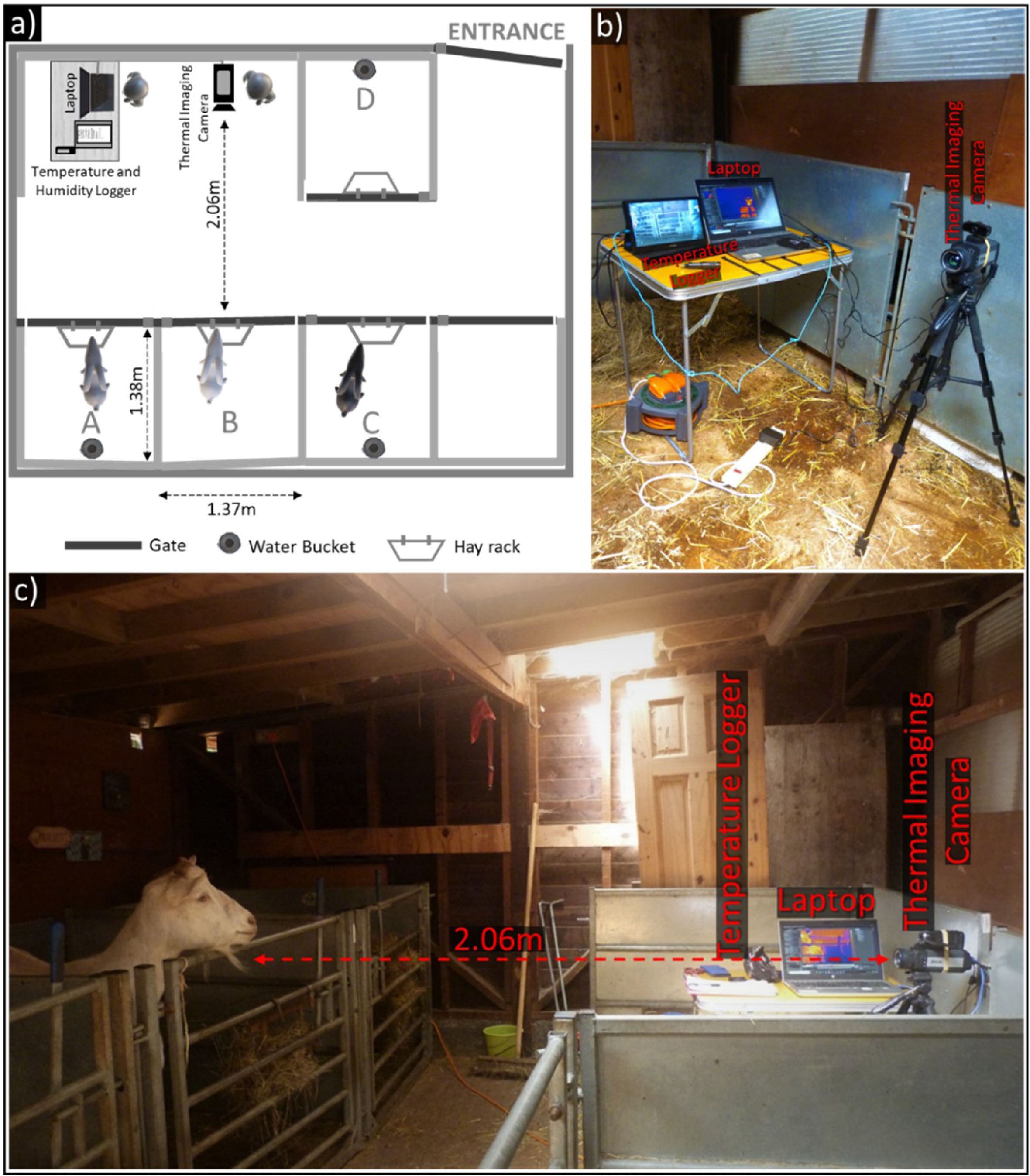
Experimental set-up for thermal measurements. a) Schematic of the stable where experiments took place. The thermal imaging camera was positioned 2.06m from the front of Pen B, where subjects were held during testing. b) Photograph demonstrating the set-up of the thermal imaging camera, laptop, and temperature logger. c) View of experimental set-up indicating relative locations of the subject and thermal imaging camera, along with other equipment.

The thermal imaging camera, FLIR A655sc (with a 25° lens), was mounted on a tripod and placed directly in front of the subject in pen B, with its height adjusted according to the subject’s head. This mid-to high-end camera has a spectral range of between 7.5–14 µm, a resolution of 640 × 480 pixels, a < 0.03°C thermal sensitivity, and ± 2°C accuracy. During trials, the camera worked in tandem with the software FLIR ResearchIR Max v. 4.40.9.30 to record thermal imaging videos and save them to an external laptop (HP ProBook 650 G4) connected via a USB and Ethernet cable. Goats tended to spend more time near the front of their pen, and the camera was placed at a distance of 2.06m away from the front of pen B. This subject-camera distance of approximately 2m is outside the recommended range for thermal imaging studies (≈ 1m, e.g., Okada et al., 2013; Church et al., 2014). However, due to the thermal imaging camera’s narrow field of view (25° × 19°, 31° diagonal), a further distance was necessary to decrease instances of the subject going out of frame and the camera position and angle having to be readjusted.

### 2.4. Experimental Preparation & Testing Procedure

We tested goats in seven groups of three individuals each. Each group was tested in one session per day over five consecutive weekdays, with the exception of one group whose final trial was carried out on Monday of the following week. Two to three groups of goats were tested per week over three weeks, with the order in which we tested groups on any given day being randomised, as was the order in which goats were tested within groups.

Before the onset of testing, a group of goats was led one by one into the stable. Each goat was placed in a separate pen, with the goat to be tested in pen B. Once the last group member had been placed in their designated pen, we waited 10 minutes before testing to minimise the effects of physical activity, human disturbance, and direct sunlight exposure on temperature readings, as well as giving animals the opportunity to habituate to the test set-up. After 10 minutes had passed, we filmed the subject’s face continuously for four minutes. Mistakes made during recording meant that although most trials were recorded at a rate of 6.25 FPS, 12 out of 100 trials had a higher frame rate (maximum 24.97 FPS). Throughout a four-minute trial, the first experimenter silently maintained the goat’s attention (without food or tactile reinforcement) while the second experimenter manually focused the thermal imaging camera. If a subject moved out of frame during their trial, we quickly adjusted the camera’s position or height and, if necessary, its angle accordingly. After four minutes had passed, the subject was moved to an adjacent pen, and the next goat to be tested was moved into pen B. To minimise the effect of disturbance and movement artefacts on temperature measures, once the goats had been rearranged, we waited five minutes before we began the next trial. Once all three subjects had been tested, they were released from the stable.

### 2.5. Video Processing & Image Analysis

A single coder extracted temperature data from the thermal imaging videos using ResearchIR. Firstly, as suitable frames were not always available (e.g., the goat was not looking at the camera), we broke each four-minute video into approximately 12s segments and one still image was extracted for each of the five consecutive segments. This measurement period generally took place within (approximately) the first 60 seconds of an experimental trial, but was shifted until later if we could not find suitable frames. A frame was considered suitable if all ROIs (left eye, right eye, and nose) were fully visible, the image was in focus, and the subject’s head was oriented at the camera, with images where the snout exceeded a 45° angle from the nasal plane being excluded from analysis. Furthermore, as the pattern of inhalation and exhalation affects temperatures in the nasal region, especially when it was possible to visualise a goat’s breathing cycle, we prioritised extracting frames just prior to the onset of inhalation (where nasal temperature began to drop).

Once five frames had been extracted per video trial, each image was calibrated according to the ambient temperature and humidity at the time of its collection. We obtained this environmental data using a EasyLog^TM^ USB-2-LC Humidity and Temperature Data Logger which recorded ambient temperature and humidity in the stable at 10s intervals during testing (to the nearest 0.5°C; mean temperature ± SD = 25.22°C ± 3.02, range = 21-31.5°C; mean relative humidity ± SD = 62.50% ± 7.17, range = 41.5-75.5%). The emissivity of the image was set to 0.98, a value generally considered to reflect that of biological tissues (Steketee, 1973), while the distance was specified as 2.06m and the reflective temperature, 20°C. Once an image had been appropriately calibrated, we used the elliptical tool to manually position ROIs. To minimise noise in temperature measures, ellipses were drawn tightly around the eye, incorporating only a thin, hairless border around each (see Figure 2). For the nose, only the tip was included, positioned in-between both nostrils. Once the ROIs had been identified, we extracted the mean, maximum, and minimum temperature from each ROI. If part of the ROI was obscured, it was not included in the analysis.

**Figure 2.**
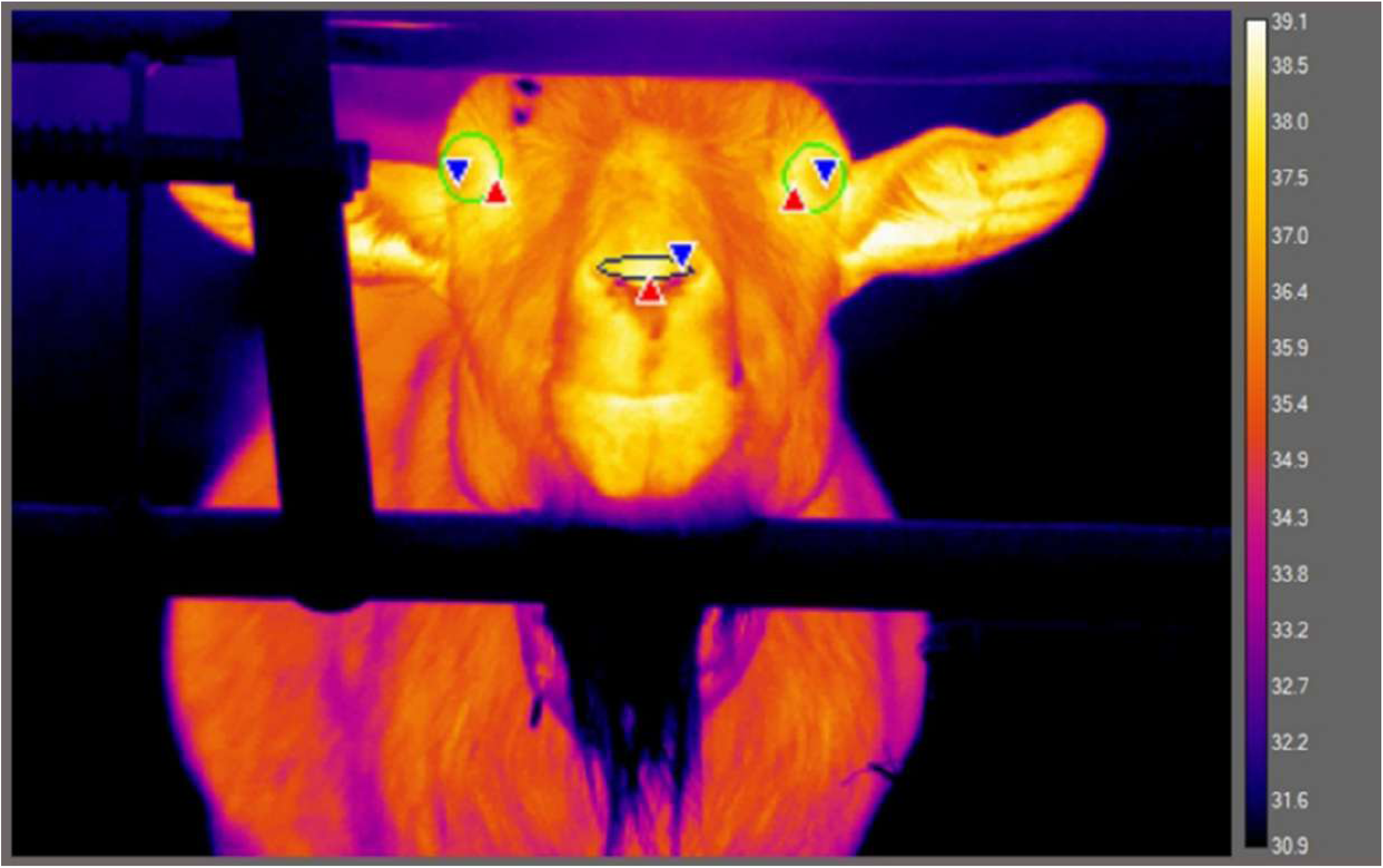
Example thermal image showing regions of interest: left eye, right eye, and nose tip (colour palette: “Fusion”). The red and blue triangles indicate the position of the pixel with the maximum and minimum temperature value, respectively, for each ROI.

To summarise, goats underwent five trials each, with five frames analysed per trial. We aimed to extract nine temperature measurements per frame, the mean, maximum, and minimum temperatures from each eye and the nose tip. Of the 21 goats tested, one was excluded for consistently not looking towards the thermal imaging camera during tests. Removing 46 instances where suitable frames could not be found, we included a total of 454 thermal images in the analysis.

### 2.6. Statistical Analysis

#### 2.6.1. Repeatability of Goat Surface Temperatures Within a Single Measurement Session

We carried out all analyses using R version 4.2.1 (R Core Team, 2022), and these were based on analyses conducted by Byrne et al. (2017). Using the package rptR (Stoffel et al., 2017) we conducted intraclass correlation analyses (ICC) to investigate repeatability of temperature measurements taken for a particular combination of ROI (left eye, right eye and nose) and measure (mean, minimum or maximum) from a single goat, in a short time frame (approximately one minute) and therefore, under relatively consistent conditions. The temperature data used in these analyses were collected during each goat’s final measurement session on the fifth day of testing. We fitted random effects models, specifying individual identity as a random effect (temperature ∼ 1 + (1| goat ID)). Models were used to partition variance explained by temperature differences between goats, which was compared against total variance, to calculate repeatability (R):

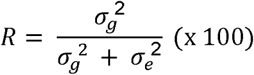

where σ_*g*_^2^ refers to the between-goat variance and σ_*e*_^2^, the error variance. R was multiplied by 100 to calculate the percentage variance explained by between-goat differences in temperature measures. 95% confidence intervals were computed around R using 10,000 bootstrapping iterations, and we employed likelihood ratio tests to investigate whether the addition of goat identity as a random effect improved model fit relative to null models excluding it.

We used the following formula to calculate the coefficient of variation (CV), for each combination of ROI and temperature measure:

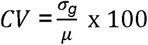

where σ_*g*_ was the standard deviation of between-goat differences in temperature, and μ was the overall mean of temperature measurements taken from the particular ROI and descriptive measure in question.

Residuals of all random effects models were visually inspected, and in combination with the DHARMa package, we evaluated how closely they conformed to assumptions of normality (Hartig, 2021). Error structures for temperature measurements taken from both eyes were approximately normally distributed, except for the mean and maximum temperatures for the right eye and the maximum temperature of the left eye. To address these deviations, we investigated the presence of influential observations or outliers using Cook’s Distance (D) in each model (*car* package: Fox & Weisberg, 2019). To meet assumptions of normality, we excluded two observations exceeding eight times the average Cook’s D, four observations exceeding five times the average Cook’s D, and the most extreme multivariate observation for the mean and maximum temperature of the right eye and maximum temperature in the left, respectively. More extreme deviations from normality were observed in goat nasal temperature measures, specifically with respect to the assumption of homoscedasticity in model residual distribution. However, as temperature is a continuous variable, transformations were ineffective at improving normality, and because alternative error structures such as Poisson distributions are more sensitive to violations of assumptions (e.g., overdispersion), we assumed a Gaussian distribution for all analyses (Knief & Forstmeier, 2021). It must be noted, although effects of heterogeneous residual variance tend not to bias variance estimates explained by random effects (from which we calculated repeatability), these estimates may become less precise (Schielzeth et al., 2020). Therefore, repeatability measures of nasal temperatures are likely to be less reliable than those of goat eye temperatures. Although Gaussian models tend to be more robust to violations against distributional assumptions, they are still sensitive to multivariate outliers with high leverage. After plotting residuals of preliminary models, we identified and removed outliers exceeding four times the average Cook’s D (nose mean temperature: 8 observations excluded; maximum temperature: 4 observations excluded; minimum temperature: 7 observations excluded). While the exclusion of these influential outliers helped improve the conformation of error residuals to assumptions of normality, they might have increased repeatability estimates due to the removal of temperature values less in line with other observations.

#### 2.6.2. Precision in Temperature Estimates

Precision was defined using the following formula:

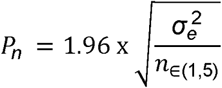

where σ_*e*_^2^ was the error variance, and *n* was the number of thermal images (1-5). The right side of this formula calculates the standard error, and altogether it estimates the 95% confidence interval range for which the mean temperatures measured from 1-5 images were expected to fall in relation to the mean of five replicate measurements (Field, 2014; Byrne et al., 2017).

#### 2.6.3. Reproducibility of Temperature Measures Taken Over Multiple Sessions

To investigate how temperature measures in each ROI varied over five days, we again used ICC analyses, this time specifying measurement session and the individual goat tested as random effects (temperature ∼ 1 + (1| measurement session) + (1| goat ID), n = 10,000 bootstraps):

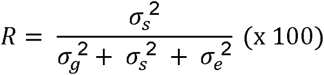

where σ_*s*_^2^ refers to the variance in temperature readings between sessions, σ_*g*_^2^ to that explained by inter-goat differences and σ_*e*_^2^, the error variance. As conditions for each measurement session were not uniform between goats (e.g., session 2 did not take place at the same time or day for each goat), we nested measurement sessions within individuals by giving every measurement session a unique ID (Schielzeth & Nakagawa, 2013). Given this nesting pattern, the effect of measurement session can be interpreted as the temperature variance explained by measurement session alone, combined with that explained by the interaction between goat identity and measurement session, or how readings from individual goats varied over the five days they were imaged (Schielzeth & Nakagawa, 2013). We used likelihood tests to investigate whether adding measurement session as a random effect significantly improved model fit against reduced models excluding this variable.

To ensure temperatures measured in the eye were approximately normally distributed, we excluded observations exceeding 10 times Cook’s D when analysing mean (6 observations excluded), maximum (2 observations excluded), and minimum temperatures of the left eye (9 observations excluded), as well as the mean of the right eye (7 observations excluded). In addition, we excluded the single most extreme observation for minimum temperatures measured in the right eye. When fitting linear random effects models to predict variance in nasal temperatures, deviations from normality were again more extreme, so a more stringent criterion of excluding observations exceeding four times Cook’s D was employed (for mean, maximum, and minimum nasal temperatures, 15, 16, and 15 outliers were excluded, respectively).

## 3. RESULTS

### 3.1. Repeatability of Goat Surface Temperatures Within a Single Measurement Session

Adding an individual-level random effect to investigate inter-goat differences improved fit of models. This helped predict surface temperature measured on each goat’s fifth day of testing for all combinations of descriptive measure and ROIs (all *p* <0.0001: Table 1). Overall, eye temperatures were restricted to a narrower range of values than those measured in the nose tip (Table 2), with the latter showing greater variability in temperature around the sample mean for each measure (higher CVs: Table 1). This variability in nasal temperatures can largely be attributed to inter-individual differences between goats, with within-goat differences over five repeated measures (error variance) being comparable to those found in eye temperatures. Consequently, repeatability of nasal temperatures was slightly higher than in either eye, but was nonetheless excellent for mean and maximum temperatures across all ROIs (rule of thumb: <0.5 poor repeatability, 0.5-0.75 moderate repeatability, 0.75-0.9 good repeatability and >0.9 excellent repeatability: Koo & Li, 2016), with between 93.50% (maximum temperature of right eye) and 99.81% (mean nasal temperature) of total variation in temperature being attributed to the individual goat tested (Table 1).

**Table 1.** Between-goat variance, within-goat error variance, coefficient of variation, proportion of variance explained by between-goat differences (repeatability: R), 95% confidence intervals for R, and *p*-values associated with the addition of between-goat differences versus null models excluding this effect for temperatures measured in each region of interest and descriptive measures on the fifth day of testing.

| Region of Interest | Measure | Individual Variance (°C<sup>2</sup>) | Error Variance (°C<sup>2</sup>) | CV (%) | R | CI | p-value |
| --- | --- | --- | --- | --- | --- | --- | --- |
| **Left Eye** | Mean | 0.440 | 0.018 | 1.815 | 0.960 | 0.915, 0.079 | <0.001 |
|  | Maximum | 0.353 | 0.019 | 1.587 | 0.949 | 0.890, 0.973 | <0.001 |
|  | Minimum | 0.968 | 0.297 | 2.817 | 0.765 | 0.569, 0.867 | <0.001 |
| **Right Eye** | Mean | 0.368 | 0.014 | 1.659 | 0.964 | 0.921, 0.981 | <0.001 |
|  | Maximum | 0.270 | 0.019 | 1.387 | 0.935 | 0.858, 0.966 | <0.001 |
|  | Minimum | 0.546 | 0.180 | 2.117 | 0.752 | 0.546, 0.859 | <0.001 |
| **Nose** | Mean | 11.612 | 0.023 | 9.691 | 0.998 | 0.996, 0.999 | <0.001 |
|  | Maximum | 6.812 | 0.019 | 7.201 | 0.997 | 0.994, 0.999 | <0.001 |
|  | Minimum | 16.498 | 0.089 | 12.027 | 0.995 | 0.988, 0.997 | <0.001 |

**Table 2.** Mean temperature, standard deviation, and temperature range for each goat facial region of interest and measures taken within a single session (day 5) and across all five sessions.

| Region of Interest | Measure | Measured Within a Single Session |  | Measured Across All Sessions |  |
| --- | --- | --- | --- | --- | --- |
| | | Mean ± SD ($^{\circ}$C) | Range ($^{\circ}$C) | Mean ± SD ($^{\circ}$C) | Range ($^{\circ}$C) |
| **Left Eye** | Mean | 36.554 ± 0.680 | 35.200 - 37.984 | 36.541 ± 0.791 | 34.226 – 38.441 |
| | Maximum | 37.443 ± 0.610 | 36.359 - 38.850 | 37.434 ± 0.687 | 35.213 – 39.257 |
| | Minimum | 34.918 ± 1.111 | 31.709 - 36.854 | 34.941 ± 1.221 | 30.678 – 37.639 |
| **Right Eye** | Mean | 36.537 ± 0.602 | 35.295 - 37.854 | 36.440 ± 0.839 | 33.660 – 38.513 |
| | Maximum | 37.457 ± 0.528 | 36.418 - 38.856 | 37.320 ± 0.726 | 34.698 – 39.150 |
| | Minimum | 34.903 ± 0.834 | 32.803 – 36.638 | 34.775 ± 1.277 | 30.497 – 37.793 |
| **Nose** | Mean | 35.163 ± 3.466 | 25.616 – 38.290 | 34.508 ± 4.041 | 21.885 – 38.691 |
| | Maximum | 36.243 ± 2.428 | 28.553 – 38.804 | 35.770 ± 3.194 | 24.828 – 39.091 |
| | Minimum | 33.772 ± 4.041 | 23.710 – 37.888 | 32.584 ± 4.985 | 20.196 – 38.158 |

With respect to descriptive measures, variability was highest in minimum temperatures (larger temperature range and higher CV), which included greater variation both between- and, moreover, within-goats, across repeated measures (Tables 1 and 2). Accordingly, minimum temperature values for the left and right eye showed lower repeatability, although it was still moderate to good, with 76.51% and 75.19% of total variation in surface temperatures, respectively, explainable by the individual goat tested (Table 1). However, for nasal temperatures, although repeatability was slightly lower (and likely negligibly so), it was still excellent, with 99.47% of temperature variance explained by differences in surface temperature between goats. By comparison, differences in repeatability between mean and maximum temperatures were more negligible across all ROIs (confidence intervals feature substantial overlap); although repeatability was slightly higher and within-goats differences, slightly lower for mean eye temperatures, with this especially being the case for the right eye. The opposite was true for nasal temperatures, with differences across repeated measures being slightly lower for maximum temperatures.

### 3.2. Precision in Temperature Estimates

The magnitude of standard error was consistently higher (i.e., precision was lower) for minimum measures, meaning more replicate thermal images would be required to achieve a similar level of precision as when using mean or maximum temperatures, across all ROIs (Table 3). For example, if we were to extract the minimum temperature of a goat’s left eye from three images, we would be expected to obtain a mean temperature within ± 0.62°C of the mean of five images 95% of the time. However, if we extracted the mean temperature in the same ROI from a single image, we would be expected to be within ± 0.26°C of the mean of five images 95% of the time. Differences in precision between mean and maximum temperatures were less, but slightly higher for goat mean eye temperatures and the maximum temperature measured in the nose tip. The ROI and measure for which we observed the highest levels of precision was in the mean temperature of the right eye (0.231-0.103°C), which also showed the least variability across repeated measurements (lowest error variance; Tables 1; 3).

**Table 3.** Precision, i.e., the standard error (°C) within which the mean of one to five replicate thermal images (*P_1-5_*) for each goat facial region of interest and measure is expected to fall in relation to the mean of five image replicates 95% of the time.

| Region of Interest | Measure | $P_1$ | $P_2$ | $P_3$ | $P_4$ | $P_5$ |
| --- | --- | --- | --- | --- | --- | --- |
| Left Eye | Mean | 0.264 | 0.187 | 0.153 | 0.132 | 0.118 |
|  | Maximum | 0.270 | 0.191 | 0.156 | 0.135 | 0.121 |
|  | Minimum | 1.068 | 0.755 | 0.617 | 0.534 | 0.478 |
| Right Eye | Mean | 0.231 | 0.163 | 0.133 | 0.115 | 0.103 |
|  | Maximum | 0.269 | 0.190 | 0.155 | 0.134 | 0.120 |
|  | Minimum | 0.832 | 0.588 | 0.480 | 0.416 | 0.372 |
| Nose | Mean | 0.294 | 0.208 | 0.170 | 0.147 | 0.132 |
|  | Maximum | 0.272 | 0.192 | 0.157 | 0.136 | 0.122 |
|  | Minimum | 0.584 | 0.413 | 0.337 | 0.292 | 0.261 |

### 3.3. Reproducibility of Temperature Measurements Taken Over Multiple Sessions

Adding the random effect ‘measurement session’ increased the fit of models predicting variation in goat surface temperatures measured over five days for all combinations of ROIs and measures (all *p* <0.0001; Table 4). The effect of measurement session (sessions 1 to 5) was strongest in the nose but still accounted for a substantial proportion of variation in eye temperatures. For instance, between-session effects explained around 80.83% and 85.85% of variance in mean and maximum nasal temperatures, respectively, and between 61.71% and 67.01% of variation in mean and maximum eye temperatures (between-session effects similar for both eyes).

**Table 4.** Between-measurement session variance, coefficient of variation, and proportion of temperature variance associated with between-session effects (repeatability: R), 95% confidence intervals for session repeatability, *p*-values associated with the addition of between-session effects against models excluding it, proportional variance associated with individual-level effects only, and that associated with within-goat error variance for each region of interest and measure.

| Region of Interest | Measure | Measurement Session |  |  |  |  | Other Effects |  |
| --- | --- | --- | --- | --- | --- | --- | --- | --- |
|  |  | Variance (°C<sup>2</sup>) | CV (%) | R | CI | p-value | Goat ID R | Proportional Error Variance |
| **Left Eye** | Mean | 0.397 | 1.724 | 0.617 | 0.429, 0.842 | <0.001 | 0.359 | 0.024 |
|  | Maximum | 0.318 | 1.506 | 0.667 | 0.477, 0.872 | <0.001 | 0.291 | 0.043 |
|  | Minimum | 0.772 | 2.515 | 0.514 | 0.354, 0.704 | <0.001 | 0.340 | 0.147 |
| **Right Eye** | Mean | 0.483 | 1.907 | 0.670 | 0.475, 0.998 | <0.001 | 0.305 | 0.025 |
|  | Maximum | 0.361 | 1.611 | 0.661 | 0.476, 0.863 | <0.001 | 0.278 | 0.061 |
|  | Minimum | 0.799 | 2.570 | 0.496 | 0.341, 0.680 | <0.001 | 0.333 | 0.171 |
| **Nose** | Mean | 14.123 | 10.891 | 0.808 | 0.612, 0.997 | <0.001 | 0.189 | 0.002 |
|  | Maximum | 8.640 | 8.216 | 0.858 | 0.665, 0.997 | <0.001 | 0.138 | 0.003 |
|  | Minimum | 19.140 | 13.426 | 0.746 | 0.546, 0.958 | <0.001 | 0.249 | 0.005 |

When considering across ROIs, minimum temperatures were more variable (larger temperature range; Table 2), including between measurement sessions, and moreover, within a session, across repeated measures taken from the same goat (Table 4). For example, within-goat variation accounted for 17.10% of total variation in minimum temperature of the right eye, but only 2.51% and 6.08% of that in mean and maximum temperatures in the same eye, respectively. Indeed, the proportion of unexplained variation was slightly larger for maximum, compared to mean temperatures across ROIs. However, although slight differences were observed between ROIs, the effect of measurement session explained a similar proportion of the total variation in goat mean and maximum surface temperatures (confidence intervals featured substantial overlap), with measurement session explaining less variation in minimum temperatures.

## 4. DISCUSSION

The use of thermal imaging in the fields of animal health and welfare research has increased over the last decade, but there remains a lack of consensus of how this technology should be best applied, including which regions and measures should be favoured when quantifying animal surface temperatures (reviews: Mota-Rojas et al., 2021; McManus et al., 2022). To address this knowledge gap, we used goats to investigate repeatability and precision of mean, maximum, and minimum eye and nasal temperatures taken from five thermal images collected in quick succession from videos (at 12-second intervals for approximately one minute). We also investigated the reproducibility of temperature estimates across five measurement sessions taking place over consecutive days. From these measurements, we assessed which combinations of ROIs and temperature measures are the most reliable for quantifying differences in goat surface temperatures. Our results indicate that although replicate measurements taken from individual goats in the short term showed substantial repeatability and high precision, surface temperatures from individual goats were not readily reproducible across days.

### 4.1. Repeatability of Goat Surface Temperatures Within a Single Measurement Session

Repeatability of five thermal images taken in quick succession was excellent for mean and maximum temperature across all ROIs, with between 93.50% (maximum temperature of right eye) and 99.81% (mean nasal temperature) of variation being attributed to the individual goat. By contrast, minimum temperatures were less repeatable and more variable both between goats and, moreover, within goats across repeated measures. As minimum temperature is derived from the value of a single pixel within a specified ROI, it is thought to be particularly vulnerable to measurement errors, the positioning of the ROI, and presence of debris or moisture on the animal’s surface (Metzner et al., 2014; Byrne et al., 2017; Uddin et al., 2020). Ultimately, given the greater variability and lower repeatability of minimum temperatures, and in line with recommendations made by similar investigations (cf. Metzner et al., 2014; Byrne et al., 2017; Uddin et al., 2020) we advocate against use of minimum temperature measures when carrying out thermal imaging in goats, and more generally, in other animals.

Given the high proportion of variation in mean and maximum temperatures explainable by between-compared to within-goat differences across repeated measures, both appear appropriate for quantifying eye and nasal temperatures over a short period. However, temperature variation across repeated measures was slightly lower, and precision was higher for mean over maximum eye temperatures (and especially in the right eye). This could suggest using the mean is more reliable for measuring eye temperatures, and as for nasal temperatures, given the slightly lower within-goats differences and higher precision observed in maximum temperatures, the latter may be more suitable. As mean temperatures are estimated from all pixels within a ROI, one key consideration affecting repeatability is the consistency with which ROI boundaries are defined across thermal images (e.g., the ROI’s size, shape, and position; Cuthbertson et al., 2019). By contrast, measuring maximum eye temperatures can often be achieved by extracting the maximum temperature of thermal images (e.g., Jerem et al., 2015), substantially reducing processing time. This is especially important for video analysis where a large number of measurement frames can be gathered in a short time (Hoffmann et al., 2013; Cuthbertson et al., 2019). However, maximum, like minimum temperatures, are calculated from the value of a single pixel, so they are more sensitive to measurement inaccuracies than the mean. According to manufacturer specifications, temperatures measured by the camera used for the current investigation are expected to be within 2°C of an object’s genuine surface temperature. These issues, to an extent, can be overcome by ensuring the highest quality thermal images, camera calibration to ambient conditions (Kim & Cho, 2021), and/ or through using data processing techniques to smooth raw temperature measures and reduce the influence of outliers (e.g., Cuthbertson et al., 2019 using video footage). Although the current research can make recommendations regarding the suitability of mean and maximum temperatures, we found negligible differences in repeatability between these measures. Researchers must carefully weigh the advantages and disadvantages of each when deciding which measure best suits their needs.

When comparing between ROIs, we found that between-goat differences against total variation were greater for nasal temperatures than for either eye. However, whether observed differences in repeatability are genuine is difficult to verify, given differences in the suitability of methods used to process and analyse nasal and eye temperature data. Specifically, the repeatability estimates for temperatures of the nose tip may be less precise than those measured in the eyes (Schielzeth et al., 2020; Knief & Forstmeier, 2021), and the more stringent removal of outliers in the former likely boosted repeatability. What can be concluded, however, is that nasal temperatures were far more variable between goats, with eye temperatures being restricted to a narrower range of values.

### 4.2. Precision in Temperature Estimates

We found that minimum temperatures measured from one image can differ by as much as 1.07°C from our gold standard, i.e., the average of five images, limiting its efficacy for making even broad distinctions, such as between stressed versus unstressed, or sick versus healthy goats. To achieve a similar precision in minimum temperatures as was found for mean and maximum temperatures, more images would be required. By contrast, a single image may be sufficient (although not recommended) to measure large temperature changes when using mean and maximum measures. We were able to attain a maximum precision of ± 0.10°C from five images using the mean temperature of the right eye. Given this region and measure also showed lower variability across repeated measures, this may highlight the suitability of this combination when measuring goat surface temperatures over a single session. More broadly, mean temperatures were slightly more precise when measuring goat eye temperatures, with maximum values performing slightly better for nasal temperatures. Goats have been reported to show an increase in eye temperature of 1.1°C following exposure to a stressor (Bartolomé et al., 2019) and 1-2.2°C differences in rectal temperature between febrile and non-febrile animals (Van Miert et al., 1984). Although changes in core temperature may not be perfectly mirrored in peripheral regions, in order to better detect fever, emotional experiences, and other physiological processes, uncertainty in surface temperature estimates must be reduced. Through averaging temperature across multiple thermal images collected in quick succession (assuming temperature is approximately stable over time), standard error will decrease proportionally with an increasing number of measurement frames, thereby increasing precision (Byrne et al., 2017). Obtaining a large number of measurement frames in a short time window is feasible when collecting thermal imaging videos (Hoffmann et al., 2013; Cuthbertson et al., 2019), but device memory and RAM capacity, as well as time needed to process video frames, can still limit the number of replicate measures it is practical to take. Nonetheless, the observed precision gained through measuring mean and maximum temperatures repeatedly over multiple thermal images may enable future researchers to effectively measure subtle surface temperature changes associated with less arousing emotional experiences (Proctor & Carder, 2015; Tamioso et al., 2017) and earlier stages of diseases in goats.

### 4.3. Reproducibility of Temperature Measurements Taken Over Multiple Sessions

We investigated the reproducibility of goat surface temperatures measured over five sessions taking place on consecutive days. Variability was always greatest (hence, reproducibility lowest) in minimum temperatures across all regions measured, including variation from unknown sources not explained by between-goat differences or session effects (e.g., measurement errors and variation in ROI placement across thermal images). Altogether, our results bring into question the suitability of minimum temperatures when using thermal imaging, both in the short- and longer-term.

Unlike eye temperatures, which were restricted to a narrower range, goat surface temperatures measured in the nose tip were highly variable. When measurements were taken across five days, a good proportion of variation in nasal temperatures could be attributed to between-session effects (74.61-85.85%), i.e., the effect of measurement session alone and how temperatures from individual goats changed across days. If not due to the more stringent removal of outliers in nasal temperature data, given the significance of measurement session, it appears surface temperatures in this region might be highly sensitive to ambient conditions specific to the time of imaging (e.g., temperature, humidity, emotional arousal and goat position: Church et al., 2014; Proctor & Carder, 2015; 2016; Ijichi et al., 2020). By contrast, given the proximity of the orbital region to the brain and its ample blood supply, eye temperatures often have a stronger association with core body temperatures compared to other peripheral regions (George et al., 2014; de Ruediger et al., 2018; Bleul et al., 2021; Kim & Cho, 2021), including in goats (preliminary study, eye-rectal temperature: *r* = 0.956; Marques et al., 2021). Nonetheless, although lower than for nasal temperatures, we found a substantial proportion of variation in eye temperatures could be attributed to differences in surface temperatures between sessions (between 49.59-67.01%). Some of this variation could be associated with shifts in core temperature (in relation to, e.g., circadian rhythm: Giannetto et al., 2020; emotional arousal: Beausoleil et al., 2004; Lees et al., 2020), but it is likely that eye temperatures were, like nasal temperatures, strongly influenced by imaging conditions.

The high variability in surface temperatures between goats and apparent sensitivity to imaging conditions observed in the nose tip, especially, likely limits the effectiveness of this ROI for detecting meaningful differences between goats (i.e., larger temperature differences, or sample sizes will be required to observe differences between treatment groups: Ledolter & Kardon, 2020). However, pre-existing variation in peripheral temperatures inherent to a sample of goats can, to an extent, be controlled for by focusing measures at the individual level. Indeed, for such a purpose, the sensitivity of a particular region to various external and internal parameters can be an asset. For example, temperatures in a chicken’s comb and wattle (which play key roles in thermoregulation), unlike eye temperatures, changed with stressor intensity, enabling finer-grained measurements of emotional responses (Herborn et al., 2015). Similarly, in ewes, more pronounced changes in temperature were observed in the muzzle relative to the eyes, so the former was considered a more practical region for detecting ovulation (de Freitas et al., 2018). As well as across repeated measurements taken from a single subject, temperatures measured in the eye, especially, have been used to compare among groups of animals, in relation to, for example, exogenous factors, like breed and sex (Jansson et al., 2021), and screening febrile from non-febrile animals (e.g., Schaefer et al., 2007; de Diego et al., 2013; Bleul et al., 2021). However, given the importance of imaging conditions on the eye, as well as nasal temperatures, it suggests such comparisons should be made with caution.

Our results suggest a high repeatability in surface temperatures measured from a single subject in the short-term (within one session) where conditions were consistent, so intuitively to compare among goats it may be better to test multiple animals under similar conditions (e.g., through testing subjects in quick succession) and preferably over multiple sessions (Uddin et al., 2020). Tighter control over environmental conditions should not only be recommended for between-subjects designs, with one investigation finding that an animal’s baseline temperature influenced the magnitude of subsequent skin temperature changes following exposure to a stressor (Herborn et al., 2015). For practical reasons, we imaged goats at a distance of just over two metres and at 90° to the nasal plane (frontal view). However, as the distance between the camera and the target object increases, intervening gases absorb a greater proportion of the radiant heat emitted by that object, so less is detected (Okada et al., 2013). Because goats were imaged at a distance exceeding that widely recommended for thermal imaging research (≈ 1m), and at a probably less than optimal angle (recommended angle: 90° to the sagittal plane), measured surface temperatures would likely have been lower than actual temperatures, and less accurate (Okada et al., 2013; Church et al., 2014; Jorquera-Chavez et al., 2019; Ijichi et al., 2020). Moreover, as subjects were free to move, the exact distance and angle between camera and subject varied between goats and across repeated measurements, introducing a less systematic source of variation into temperature estimates. Additional considerations include how best to define eye temperature, in particular. In the eye, the posterior border of the eyelid and lacrimal caruncle especially are known to be richly supplied by a dense array of capillary beds (Kim & Cho, 2021; Mota-Rojas et al., 2021). Indeed, investigations subsetting the eye into multiple orbital regions have found ROIs associated with specific areas (e.g., medial canthus and lacrimal sac) correlate better with rectal temperature than whole-eye measurements (Kim & Cho, 2021; Shu et al., 2022). Ultimately, through exerting greater control over the environmental component of animal surface temperatures, future investigations may be able to achieve greater reproducibility in temperature estimates (Church et al., 2014).

## 5. CONCLUSION

We found that mean and maximum surface temperatures measured in the eyes and nose tip in goats to be highly repeatable, at least in the short term. In addition, these temperature measures showed high levels of precision, with a single image potentially being enough to make broad distinctions, such as sick from healthy, or stressed from unstressed animals; although using more than one image is recommended to enhance precision. However, given the strong influence of measurement sessions, goat surface temperatures were not readily comparable across days, highlighting the importance of ambient imaging conditions on temperature estimates. Researchers using thermal imaging in small ruminants should consider focusing measurements at the individual level, and/or further refining the methodology used here (e.g., using a more optimal measurement distance and angle), as well as exerting tighter control over ambient conditions and perhaps using a more precise ROI (e.g., localised to a specific orbital region). Given the non-invasive nature of thermal imaging and the importance of animal body surface temperatures as an indicator of animal health and welfare, investigations like ours are becoming increasingly important to identify approaches in which to effectively exploit this technology to its fullest potential.

## Acknowledgements

We thank Prof. Stuart Semple and Dr Leanne Proops for their helpful advice on the research. We acknowledge the invaluable assistance of Daniela Bernal Vega and Ellen Ye, among others, during testing, and the staff and volunteers at the Buttercups Sanctuary for Goats (https://www.buttercups.org.uk/) for help and advice, as well as for access to their animals.

## Funding

This work was part-supported by a research grant from Farm Sanctuary to A.G.M. for the purchase of a thermal imaging camera, with other expenses covered by a grant from Ede & Ravenscroft to M.A.M. and the University of Roehampton. The funders had no role in study design, data collection and analyses, decision to publish, or preparation of the manuscript.

## Competing Interests

The authors declare that they have no competing interests.

## Author Contributions

- Debottam Bhattacharjee analysed the data, prepared figures and/or tables, authored or reviewed drafts of the paper, and approved the final draft.
- Marianne Mason conceived and designed the experiments, performed the experiments, collected data, analysed the data, authored or reviewed drafts of the paper, and approved the final draft.
- Alan McElligott conceived and designed the experiments, authored or reviewed drafts of the paper, and approved the final draft.

## Animal Ethics

All animal care and testing protocols were in line with ASAB guidelines for the use of animals in research (Bee et al., 2021). Approval was granted through an ethical amendment (03.21) made to the project ‘goat perception of human cues’ (Ref. LSC 19/ 280) by the University of Roehampton’s Life Sciences Ethics Committee. All procedures were non-invasive, goats were kept unrestrained and tested in groups to avoid social isolation, showing no obvious signs of stress during experimental trials.

## Data Availability

Raw data and R script will be made available upon publication.

## Notes

### Competing Interest Statement

The authors have declared no competing interest.

